# Habitat preference predicts genetic diversity and population divergence in Amazonian birds

**DOI:** 10.1101/085126

**Authors:** Michael G. Harvey, Alexandre Aleixo, Camila C. Ribas, Robb T. Brumfield

**Affiliations:** Department of Biological Sciences and Museum of Natural Science, Louisiana State University, Baton Rouge, Louisiana, 70803, USA; Department of Ecology and Evolutionary Biology and Museum of Zoology, University of Michigan, Ann Arbor, Michigan, 48105, USA; Museu Paraense Emílio Goeldi (MPEG), Caixa Postal 399, Belém, PA 66040-170, Brazil; Instituto Nacional de Pesquisas da Amazônia (INPA), Av. André Araújo 2936, Manaus, AM 69060-001, Brazil

**Keywords:** population genetics, phylogeography, habitat selection, ultraconserved elements, trait-dependent diversification, Amazon rainforest

## Abstract

The ecological traits of organisms may predict important evolutionary parameters such as genetic diversity, population genetic structure, and demographic history. Making these ecological-evolutionary links is difficult because robust, comparable genetic estimates are required from many species with differing ecologies. In Amazonian birds, differences in habitat preference are an important component of ecological diversity. A subset of Amazonian birds is restricted to forest edge and open forest along floodplains, whereas another subset occurs only in the interior of tall, upland forest. Here, we examine the link between habitat and evolutionary metrics using 20 pairs of closely related and co-distributed bird species in which one member of the pair occurs primarily in forest edge and floodplains, and the other occurs in upland forest interior. We use standardized geographic sampling and genomic data from the same set of 2,416 independent markers to estimate genetic diversity, population structure, and demographic history in each species. We find that species of upland forest have higher genetic diversity, greater divergence across the landscape, more genetically distinct populations, and deeper gene histories than floodplain species. Our results reveal that species ecology in the form of habitat preference is an important predictor of genetic diversity and divergence and suggest that floodplain and upland avifaunas in the Amazon may be on separate evolutionary trajectories and require different conservation strategies.

## Introduction

Genetic and phenotypic variation within species determines how they respond to environmental change (Willi et al. 2006), their propensity to form new species (Riginos et al. 2014), and their susceptibility to extinction (Keller and Waller 2002). Levels and geographic patterns of variation differ widely among species (Taberlet et al. 1998; Soltis et al. 2006), in many cases because they have been subject to different histories of landscape change (Lorenzen et al. 2012). In co-distributed species that have evolved under similar landscape histories, however, we may have to invoke other factors to explain differences in variation (Lessios 2008). Although stochasticity in evolutionary history may account for some differences, variation in ecology and life history among species may have additional, deterministic effects on their evolutionary trajectories.

The importance of organismal traits in determining the standing genetic diversity observed within species has received attention because of interest in the adaptive and evolutionary potential of levels of genetic polymorphism and mutation rates (Nevo et al. 1984; Leffler et al. 2012; Romiguier et al. 2014; Miraldo et al. 2016). Genetic divergence between populations is also of interest due to its potential evolutionary importance - divergent populations represent potential incipient species. Although few ecological traits have been examined, studies have found that population divergence is predicted by growth form, breeding system, floral morphology, pollination mechanism, seed dispersal mode, phenology, life cycle, and successional stage in woody plants (Loveless and Hamrick 1984; Duminil et al. 2007; Gianoli et al. 2016); microhabitat preference (branch circumference) and elevation in Costa Rican orchids (Kisel et al. 2012); larval dispersal mode in a variety of marine organisms (Palumbi 2003; Hellberg 2009); a preference for forest canopy or understory in Neotropical birds (Burney and Brumfield 2009); and body size and reproductive mode in frogs (Pabijan et al. 2012; Paz et al. 2015). However, most studies of trait-dependent divergence have been limited to estimates based on a single locus, and estimates of parameters aside from diversity and divergence have scarcely been investigated (Papadopoulou and Knowles 2016).

Genome-wide approaches to genetic sampling can provide improved estimates of genetic diversity, population genetic divergence, and other evolutionary metrics. Methods for sequencing reduced representation libraries of genomic DNA can be used to obtain information from many independent parts of the genome and many samples (e.g., Davey et al. 2011; Faircloth et al. 2012). Increasing the number of loci under investigation provides more precise estimates of parameter values that are less subject to biases resulting from coalescent stochasticity (Edwards and Beerli 2000; Carling and Brumfield 2007). Sampling hundreds of loci is equivalent to sampling an entire population at a few loci, and with enough loci many parameters can be reliably estimated even when populations are represented by only a single diploid individual (Willing et al. 2012). Datasets with many independent loci may provide sufficient power to evaluate parameter-rich models of population history that include estimates of migration between populations, demographic changes, and selection in addition to divergence (Carstens et al. 2013). Finally, processes like admixture and selection may be evident only in subsets of the genome (Counterman et al. 2004; Wall et al. 2009), and can only be detected with dense genomic sampling. Genome-wide data therefore have the potential to provide more precise and complete estimates of genetic metrics for comparison with species trait information.

The avifauna of the Amazon Basin in northern South America provides an excellent system in which to investigate the effect of traits on genomic diversity and population history. The Amazonian avifauna is the most diverse in the world (Pearson 1977) and comprises species with a variety of ecological traits (Parker et al. 1996) and, based on the few species with data, differing levels of genetic variation (Bates 2000, Smith et al. 2014b). Many species are habitat specialists (Kratter 1997, Rosenberg 1990, Alonso et al. 2013) and closely related species often partition space by associating with different habitats. Two habitats in particular, floodplain forest along whitewater rivers (*várzea*) and upland forest (*terra firme*), are widespread and are inhabited by a suite of pairs of closely related species that segregate by habitat (Remsen and Parker 1983) and sometimes exhibit interspecific aggression (Robinson and Terborgh 1995). Floodplain forest has an open, edge-like structure as a result of disturbance during floods (Prance 1979; Wittmann et al. 2004), and many floodplain species occur outside of floodplains in other edge habitats such as the borders of savanna or human-made clearings. Upland forest, conversely, is typified by a high proportion of tall trees, a dark interior, and open understory (Campbell et al. 1986; Gentry and Emmons 1987), and many upland forest species avoid open areas. It remains unclear whether Amazonian birds in floodplains and edge habitats differ from those in upland forest in genetic diversity, divergence across the landscape, or other aspects of evolutionary history.

In this study, we test whether habitat preference in Amazonian birds predicts genetic metrics of diversity, divergence, and history. We examine 40 species or species complexes (all of which are hereafter referred to as “species” for brevity) of broadly co-distributed Amazonian birds that differ in habitat association. The forty species include twenty pairs in which one species is found in upland forest, and the other is a closely related species found in floodplains and edge habitats. We use genomic sequence data from populations randomly distributed across the Amazon to estimate genetic diversity, population genetic structure, and demographic history in each species for comparative analysis.

## Methods

### Sample Design

We designed a sampling strategy to minimize the potential effects of sampling bias across species on comparisons of genetic metrics. Using published data (Parker et al. 1996; del Hoyo et al. 2002-2011; Schulenberg et al. 2010; Remsen et al. 2015) and expert knowledge (B. M. Whitney and L. N. Naka, pers. comm.), we selected genera that contained a pair of species or species complexes that generally segregate between floodplains and upland forest. Some of the genera have since been split into multiple genera (Remsen et al. 2015), but the species selected are still closely related (<0.5% average genetic distance; fig. 1). Species pairs are not sister taxa, as all are more closely related to populations or species outside of the Amazon Basin. We obtained lists of vouchered tissue samples collected during our fieldwork and available from existing natural history collections. From an initial list of 57 pairs that fit our criteria, we removed any pair containing a species for which fewer than 20 tissue samples were available in existing museum collections. The result was a list of 20 species pairs from 15 avian families (fig. 1).

**Figure 1:**
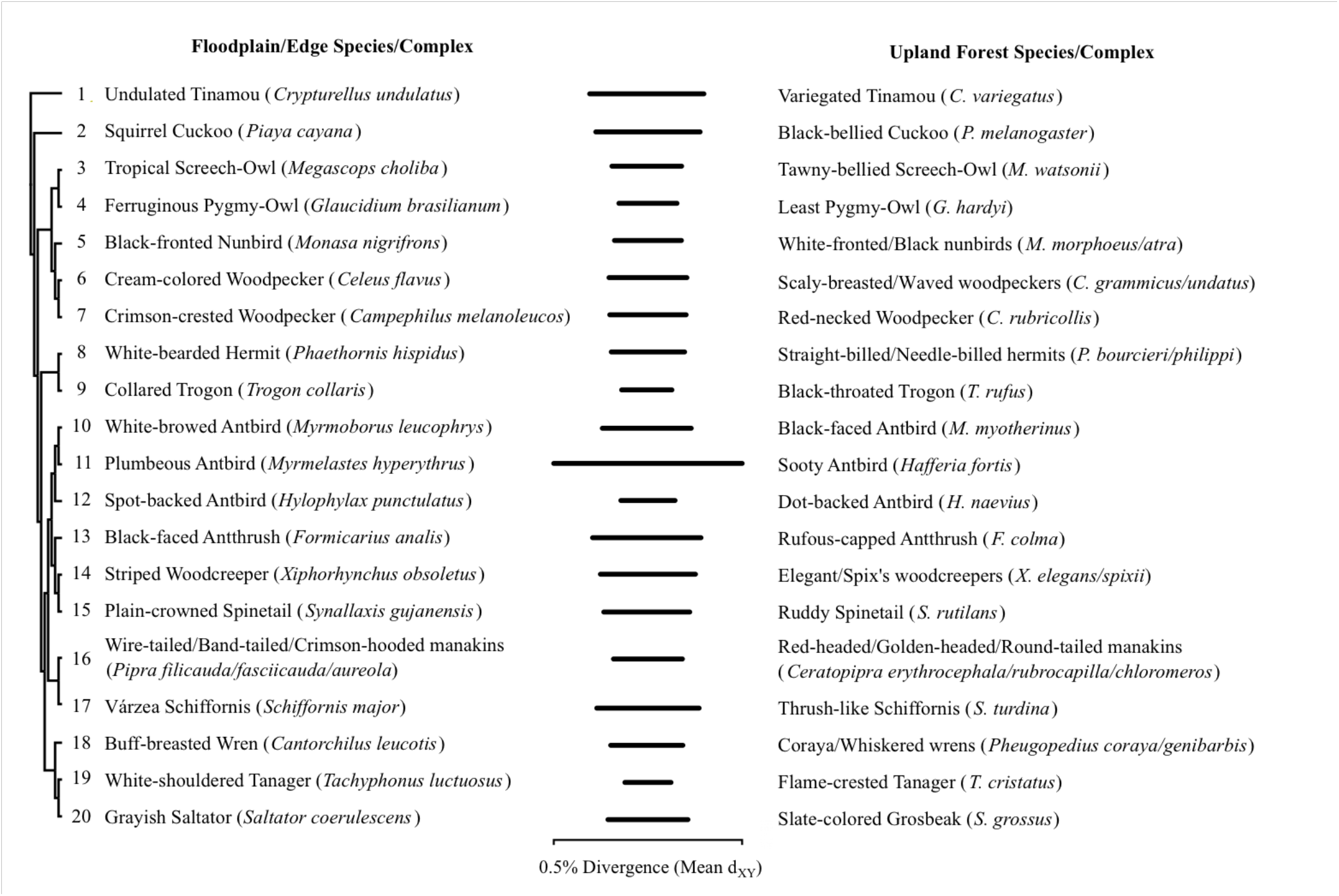
Pairs of study species or species complexes examined. A phylogenetic tree based on a MrBayes analysis of concatenated sequences depicts the relationships among the pairs, and horizontal bars in the middle of the figure depict the mean pairwise sequence divergence (*d*_XY_) between members of a pair.

For each species, we examined all populations within the Amazon. We did not include populations that appear to replace study species geographically based on distributional information, but are distantly related and would result in paraphyly of the study populations. For example, Chestnut-rumped Woodcreeper (*Xiphorhynchus pardalotus*) appears to replace our study species Elegant Woodcreeper (*X. elegans*) in the Guianas, but is in fact phylogenetically related to Ocellated Woodcreeper (*X. ocellatus*)(Sousa-Neves et al. 2013). Some study species have populations outside of the Amazon Basin, generally in the Atlantic Forest of southeastern South America or the humid forests of Central America and the Chocó region of northwestern South America, but for comparability we examined only Amazonian populations.

We selected a set of samples for each species that would minimize differences in the spatial dispersion of samples across species. We first georeferenced all genetic samples with locality information more precise than department or state and sufficient precision to determine on which side of any major biogeogeographic barriers (rivers or mountains) the sample originated. Locality records were plotted using ArcMap 10.0 (ESRI, Redlands, CA) with the WGS84 projection. We also digitized the Amazon terrestrial areas of endemism based on da Silva et al. (2005). We plotted 40 random points across the Amazon using the genrandompts function in Geospatial Modelling Environment v. 0.7.1.0 (Spatial Ecology LLC, Toronto, Canada), with a minimum distance between points of 2 map units (equivalent to two degrees in WGS84) and requiring 2 or more points within each area of endemism (da Silva et al. 2005). For each species, we determined the closest sampling locality to each random point using the spatial join function in ArcMap. Due to the vagaries of sample availability, some samples were quite clustered. We thinned sampling to 20 individuals per species by projecting the sample localities on a blank grid and removing clustered samples without reference to the underlying geography.

### Laboratory Methods

We extracted whole genomic DNA from tissue samples using DNeasy Blood and Tissue kits (Qiagen, Valencia, CA) and quantified extracts using a QuBit fluorometer (ThermoFisher, Waltham, MA). We excluded samples with extracts containing less than 1 µg of DNA total. We thinned sampling based on spatial dispersion without reference to geography, as described above, to arrive at a final set of 11 samples for each species.

Due to the comparative nature of our study, it was important to obtain genetic data that would not bias estimates of genetic diversity and population history across species. Results are generally not comparable across species if different loci are examined, because orthology assessment among sequence reads leads to biased levels of variation (Harvey et al. 2015). Sequence capture of conserved genomic regions permits the interrogation of the same loci across divergent species (Faircloth et al. 2012; Bi et al. 2012; Hedtke et al. 2013), and orthology assessment in the assembly of sequence capture datasets is straightforward and has relatively little impact on allelic diversity (Harvey et al. 2016).

We used sequence capture to target ultraconserved elements (UCEs) and exons from across the genome. We modified existing sequence capture probe sets for UCEs (Faircloth et al. 2012) to obtain additional sequence from the more variable UCE flanking regions that might be useful for inferring shallow population histories. In UCE loci targeted with a single probe, we designed two probes extending further into the UCE flanks. The 120-mer probes were tiled such that they had 50% overlap (60 bp) in the middle of the locus and covered 180 bp total. Probe sequences were based on the chicken (*Gallus gallus*) genome release ICGSC Gallus_gallus-4.0 (Hillier et al. 2004). We also targeted conserved exons adjoining variable introns that have been used in previous avian phylogenetic studies (Kimball et al. 2009; Wang et al. 2012; Smith et al. 2012). Probes were designed off the chicken genome sequence and were again tiled such that they covered the entire exon sequence at 2x coverage (50% overlap between adjoining probes). The final probe set included 4,715 probes targeting 2,321 UCEs and 96 exons.

We sent all samples to Rapid Genomics (Gainesville, FL) for sequence capture and sequencing following the general protocol described in Faircloth et al. (2012) and Smith et al. (2014a). Samples were multiplexed at 160 samples per lane on a 100 bp paired-end Illumina HiSeq 2500 run. Rapid Genomics demultiplexed raw reads using custom scripts and strict barcode matching.

### Bioinformatics

We cleaned reads with Illumiprocessor (Faircloth 2013). We developed a pipeline (seqcap_pop; https://github.com/mgharvey/seqcap_pop) to process and assemble datasets as follows. We used Velvet (Zerbino and Birney 2008) and the wrapper program Velvet Optimiser (Gladman 2009) exploring hash lengths of between 67 and 71 to assemble reads across all individuals into contigs *de novo*. We mapped contigs to UCE probe sequences using Phyluce (Faircloth 2015). For each individual, we mapped reads to contigs that aligned to UCEs using bwa (Li and Durbin 2009). We explored thresholds that allowed anywhere from 1 to 7 mismatches between reads for mapping (Harvey et al. 2015) and, based on plots of allele loss, selected a setting of 4 permitted mismatches per read for final assemblies. We converted sam files to bam format using samtools (Li et al. 2009) and cleaned bam files by soft-clipping reads outside the reference contigs with PICARD (Broad Institute, Cambridge, MA). We added read groups for each individual using PICARD and merged the bam files across individuals with samtools. We realigned reads to minimize mismatched bases using the RealignerTargetCreator and realigned indels using IndelRealigner in the GATK (McKenna et al. 2010). We called single nucleotide polymorphisms (SNPs) and indels using the GATK UnifiedGenotyper, annotated SNPs with VariantAnnotator, and masked indels using VariantFiltration. We removed SNPs with a quality score below Q30 and conducted read-backed phasing using the GATK. We output SNPs in vcf format and used add_phased_snps_to_seqs_filter.py from seqcap_pop to insert SNPs into reference sequences and produce alignments for each locus across individuals. SNPs on the same locus for which phasing failed were inserted using the appropriate IUPAC ambiguity codes. We collated sequences and produced final alignments using MAFFT (Katoh et al. 2005).

We also assembled partial mitochondrial genomes for each sample from off-target reads using a similar pipeline. We obtained existing complete or nearly complete mitochondrial genome sequences from the most closely related taxon to each study species for which they were available (table A1). We mapped reads to the mitochondrial genomes, sorted the bam file, recalculated MD tags, and indexed the bam file using Samtools. We then called variant sites and output vcf files containing variant and invariant bases using Freebayes (Garrison and Marth 2012) and used these to assemble sequences using freebayes_vcf2fa_mt.py (https://github.com/mgharvey/misc_python/bin/freebayes_vcf2fa.py). Only sites with a read depth of 5 or greater were included in sequences. We conducted final alignment with MAFFT.

We searched for potential sample identification errors or signs of contamination by building exploratory trees of concatenated SNPs from the UCE/exon data using MrBayes v.3.2.2 (Ronquist et al. 2013) and scrutinizing any long branches and by mapping mitochondrial sequences to existing sequence data in Genbank (Benson et al. 2014) using Blastn (Altschul et al. 1997). We counted the reads in BWA assemblies using Samtools.

### Summary Statistics of Diversity

We calculated basic population genetic summary statistics including number of variable sites, nucleotide diversity (π)(Tajima 1983), Watterson’s θ (Watterson 1975), and Tajima’s D (Tajima 1989) across all samples in each species using DendroPy v.3.10.0 (Sukumaran and Holder 2010). We calculated average heterozygosity within individuals for each species as a measure of the standing genetic diversity within populations. We estimated gene trees for each locus in each species using RAxML v.8 (Stamatakis 2014). Genetic variation may differ among genomic regions, such as on sex-linked chromosomes versus autosomes (Counterman et al. 2004). We determined the genomic location of recovered loci by mapping them to the Zebra Finch (*Taeniopygia guttata*) genome (Warren et al. 2010). We then compared levels of nucleotide diversity on loci mapping to the Z chromosome to those mapping to the autosomes.

### Population Genetic Structure

We examined multiple strategies for estimating divergence among individuals. We first estimated simple summaries of overall population genetic structure in each species using F_ST_ and d_XY_. We estimated F_ST_ among individuals in each species using the statistic developed by Reich et al. (2009), which has been shown to be unbiased and effective even when dealing with sample sizes as small as two alleles per population (Willing et al. 2012). We estimated d_XY_ among individuals using average sequence distance between samples after correcting for multiple substitutions using the method of Jukes and Cantor (1969).

We next examined methods to infer population clustering across individuals and assign individuals to populations. Various methods are available to infer population structure, and they can produce different results (Latch et al. 2006; Chen et al. 2007). We therefore examined results from three alternative methods: Structure (Pritchard et al. 2000), Bayesian Analysis of Population Structure (BAPS; Corander et al. 2003), and Discriminant Analysis of Principal Components (DAPC; Jombart et al. 2010). We also used the first two methods to determine if any of the individuals sampled were assigned with high probabilities to multiple populations, suggestive of admixture between populations. Structure is a model-based clustering method that simultaneously infers population structure and assesses the probability of individual assignment to a cluster or combination of clusters. We ran Structure using the linkage model, and provided phase information for each site in each individual as well as distances in base-pairs between linked sites. Sites mapping to different loci were treated as unlinked. We conducted analyses at k-values ranging from 1 to 6, with 10 replicate runs at each value. Each run included a 50,000-iteration burn-in followed by 200,000 sampling iterations, and we assessed convergence by examining alpha, F, D_ij_, and the likelihood within and across runs at each k-value. We estimated the best value of k using the method of Evanno et al. (2005) implemented in StructureHarvester (Earl 2012). In some cases, the results at the best k value included clusters to which no individuals were assigned. In these situations, we also examined the largest k value in which at least one individual was assigned to each cluster. We combined results across replicates runs with the best k value using CLUMPP (Jakobsson and Rosenberg 2007).

BAPS is a model-based clustering method that jointly infers the number of populations and population assignment of individuals, which can then be used in a subsequent analysis of admixture for each individual. Because BAPS requires complete phasing information for linked sites, and phasing had failed for some individuals at most linked sites in our datasets, we used the unlinked model and examined only a single randomly selected SNP from each locus for this analysis. We conducted mixture clustering with the maximum number of populations (k) set at 10. We estimated admixture in each individual based on mixture clustering using 50 simulation iterations, 50 reference individuals, and 10 iterations to estimate admixture coefficients in the reference individuals.

DAPC is a fast, non-parametric method for inferring the number of genetic clusters and cluster assignments in large datasets. We inferred the number of clusters and cluster membership in DAPC using the maximum number of principal components available for each species, and selected the best value for cluster number by choosing the value at which Bayesian Information Criterion reached a low point (Jombart et al. 2010). Unlike Structure and BAPS, DAPC does not allow for admixture estimation.

### Demographic Modeling

We estimated demographic parameters using a coalescent modeling approach in G-PhoCS v.1.2.3 (Gronau et al. 2011). We ran analyses using all population assignments inferred in Structure, BAPS, and DAPC to assign population membership. In Structure results, individuals with multiple assignments were placed in the population with the highest assignment probability. We specified the population topologies in situations where more than two populations were present based on the MrBayes trees of concatenated SNPs. For each species, we examined both a model with no migration between populations subsequent to divergence as well as a model allowing for migration between terminal populations. We used gamma priors of (1, 5000) for θ and τ and (1,3) for migration and ran runs for a minimum of 500,000 iterations (sampling every 100). We also explored the impact of θ and τ priors of (1, 50). Convergence was assessed by examining parameter traces and ESS values in Tracer v.1.5 (Rambaut and Drummond 2007). G-PhoCS implements a multi-population model and cannot be run in the study species with a single population. For comparative analyses, we used the species-wide θ values from DendroPy and divergence time (τ) values of zero for single-population species.

### Comparative Analyses

The above analyses produced 18 metrics representing summaries of dataset attributes, genetic diversity, differences in diversity between the Z chromosome and autosomes, genetic divergence across space, and historical demography (table 1). We expect many of these metrics to exhibit correlations with each other. We estimated Spearman’s correlations between all pairs of variables and significance using the R package Hmisc (Harrell 2016), and also grouped highly correlated variables using the ClustOfVar R package following developer recommendations (Chavent et al. 2012). We examined whether each genetic metric was associated with the level of evolutionary relatedness among study species. We estimated a phylogeny for all 40 species by aligning UCE and exon sequences from one sample of each species in MAFFT. Because sequences were assembled by mapping to different contigs in each species, the sequences were generally not entirely overlapping across species, and these ragged ends frequently included messy and potentially spuriously aligned blocks of sites. We removed these by filtering for only sites without missing data in the alignments. We concatenated filtered alignments that contained all 40 individuals and conducted a Bayesian phylogenetic analysis on the complete matrix in MrBayes. We square-root transformed right-skewed genetic variables to achieve normality and calculated phylogenetic signal in each variable using Pagel’s λ in the R package Phytools (Revell 2011), with 999 permutations to assess whether λ differed significantly from zero. We also tested whether the degree to which each variable differed between members of a species pair was predicted by the overall level of pairwise sequence divergence between them. Because some of the study species contained multiple named species under current taxonomy (see Sample Design above), we tested whether this could explain variation in genetic metrics.

**Table 1:** Associations between habitat preference and genetic metrics

| Metric | Group | GLMM<br>*t* | GLMM<br>*P* | PGLS<br>*t* | PGLS<br>*P* |
| --- | --- | --- | --- | --- | --- |
| Data attributes: |  |  |  |  |  |
| Mapped Reads | 1 | .25 | .80 | 1.32 | .20 |
| Assembled Sequence Length | 2 | -.77 | .45 | -.98 | .33 |
| Average Number of Variable Sites | 3 | 3.47 | .001 | 4.92 | <.001 |
| Summary statistics of diversity: |  |  |  |  |  |
| Nucleotide Diversity ($\pi$) | 2 | 2.08 | .04 | 3.29 | .002 |
| Z:A Chromosome Nucleotide Diversity ($\pi$) | 5 | .34 | .74 | 1.08 | .29 |
| Watterson's $\theta$ | 3 | 3.36 | .002 | 4.81 | <.001 |
| Tajima's $D$ | 3 | -2.03 | .05 | -2.31 | .03 |
| Within-Population Heterozygosity | 2 | -.02 | .98 | -.43 | .67 |
| Average Gene Tree Height | 2 | 2.39 | .02 | 3.59 | <.001 |
| Population genetic structure: |  |  |  |  |  |
| Mean $d_{XY}$ Between Populations | 2 | 2.10 | .04 | 3.30 | .002 |
| Mean $F_{ST}$ Between Populations | 4 | 2.19 | .03 | 2.89 | .006 |
| Number of Structure Populations | 4 | 1.22 | .22 | 2.80 | .008 |
| Number of BAPS Populations | 4 | .62 | .54 | 1.64 | .11 |
| Number of DAPC Populations | 4 | 1.28 | .20 | 2.88 | .007 |
| Demographic model parameters (G-PhoCS): |  |  |  |  |  |
| Average $\theta$ Across Populations | 6 | .10 | .92 | -.15 | .88 |
| Change in $\theta$ Through Time | 6 | -0.23 | .82 | -.07 | .94 |
| Oldest Population Divergence or $\tau$ | 4 | .03 | .98 | 1.59 | .12 |
| Average Migration Rate | 7 | .08 | .94 | -.48 | .64 |

Note: Group numbers indicate assignment to clusters of semi-independent variables based on ClustOfVar. Results are from single-predictor tests.

We tested whether habitat predicted metrics of population genomic diversity and population history using two strategies to account for shared evolutionary history. We first used generalized linear mixed models (GLMMs) to test for correlations, using genus as a random variable to account for the shared history between study species pairs. The generalized linear modeling approach allowed us to examine response variables with diverse error distributions in the same statistical framework. Gaussian error models were used for continuous and large count data, Poisson models for data composed of low count values (<100), and Gamma models with a logarithmic link function for continuous data with positive skew. We examined the relationship between habitat and each genetic response variable in one-way tests using functions for GLMMs in the stats R package (R Core Team 2015). Covariance due to shared history can also be modeled using phylogenetic distance. We square-root transformed right-skewed data to achieve normality and used Phylogenetic generalized least squares (PGLS) in the R package caper (Orme et al. 2013) to test for associations between habitat and genetic metrics while controlling for relatedness among species with the MrBayes phylogeny of concatenated data.

Although our primary focus was on the associations between habitat and genetic diversity, we also examined two additional traits thought to predict population divergence in Neotropical birds. First, whether a bird inhabits the forest canopy or understory has been shown to predict levels of divergence across landscape barriers (Burney and Brumfield 2009, Smith et al. 2014b), so we tested whether canopy and understory species (based on Parker et al. 1996) differed in metrics of population genomic diversity. Second, habitat or microhabitat associations may affect population genetic divergence via differences in dispersal ability among species (Burney and Brumfield 2009). We examined whether Kipp’s index, a morphological index of dispersal ability that can be measured from museum specimens (Kipp 1959), predicted levels of population genomic diversity across species. Because species differing in forest stratum and Kipp’s index were not organized into pairs, we used PGLS rather than GLMMs to test for correlations with genetic variables. Finally, we tested whether associations between habitat and genetic metrics could be explained by second-order interactions using multi-predictor GLMM and PGLS models combining forest stratum and/or Kipp’s index with habitat.

## RESULTS

We obtained an average of 2,087,266 (SD = 656,446) raw reads per sample. On average, 28.1% (SD = 6.57%) of sequence reads were successfully mapped to target loci after cleaning and 0.44% (SD = 0.60%) of all reads mapped to the mitochondrion. Across species, we obtained data from an average of 2,142 UCEs (SD = 65.5) and 69 exons (SD = 4.8). We recovered data in at least one species from 2,416 of 2,417 targeted loci. Mean locus length was 554 bp (SD = 56.3), and there were 7,196 (SD = 1,379) sites that were variable, on average. Additional summary statistics are provided in table A2 and appendix B.

Based on MrBayes trees of concatenated SNPs and Blastn results of mitochondrial sequences, we determined eight samples were likely misidentified or heavily contaminated and were removed from further analyses (table A3). Three samples contained large numbers of rare alleles likely to be a result of lower levels of contamination or sequencing errors and were also removed (table A4). Removing these samples resulted in concatenated SNP trees with low to moderate structure based on internal branch lengths (fig. A1). Three samples failed, with greater than 85% missing data at variable sites, and were removed (table A5). After removing these 14 samples, we were left with 440 samples (plus 24 extra-Amazonian outgroup samples) across the 40 study species.

Nucleotide diversity averaged 1.09×10^-3^ (SD = 2.98×10^-4^), Watterson’s θ averaged 0.79 (SD = 0.22), and Tajima’s D averaged -0.79 (SD = 0.36) across all samples. Average F_ST_ was 0.26 (SD = 0.14) and average d_XY_ was 1.11×10^-3^ (SD = 3.10×10^-4^). Gene tree depths averaged 3.93×10^-3^ (SD = 7.67×10^-4^). Across study species, contigs from 2,415 of 2,416 recovered loci successfully mapped to the Zebra Finch genome assembly. Contigs from all species mapped to the Z chromosome for 171 loci, to one of the autosomes for 2,169 loci, and to unplaced scaffolds in 44 loci. For 31 loci, contigs from different species mapped to different chromosomes or scaffolds resulting in ambiguous positions. Based only on loci mapping to the Z chromosome or autosomes in all study species, the ratio of nucleotide diversity on the Z chromosome to that on the autosomes averaged 1.04 (SD = 0.263).

The number of populations and population assignments inferred from Structure, BAPS, and DAPC were broadly concordant (figs. 2, A2). The best k-value from Structure analyses based on the Evanno method, after reducing k to remove clusters without assigned individuals, ranged between one and four across study species (median = 3). The number of populations estimated in BAPS varied from one to three (median = 2), and in the number of clusters from DAPC varied between one and four (median = 2). Many individuals contained mixed probabilities of assignment to different clusters in the Structure results, potentially indicative of admixture, but no admixture was recovered in the admixture analysis from BAPS. Populations from all three methods were generally partitioned among geographic areas, with boundaries broadly concordant with major rivers (fig. A3).

Estimates of historical demography from G-PhoCS for the 23 species with multiple populations (fig. 2, appendix C) revealed an average per-site θ across populations of 1.53×10^-3^ (SD = 5.73×10^-4^). θ values in contemporary populations averaged 2.68 times larger than the θ inferred for the ancestral population at the root (SD = 1.29). The height of the deepest divergence in the model (τ) varied from 9.72×10^-5^ to 1.13×10^-3^ across species (mean = 4.45×10^-4^). Average migration rate between populations within a species varied from 0.337 to 4.69 (mean = 0.950).

**Figure 2:**
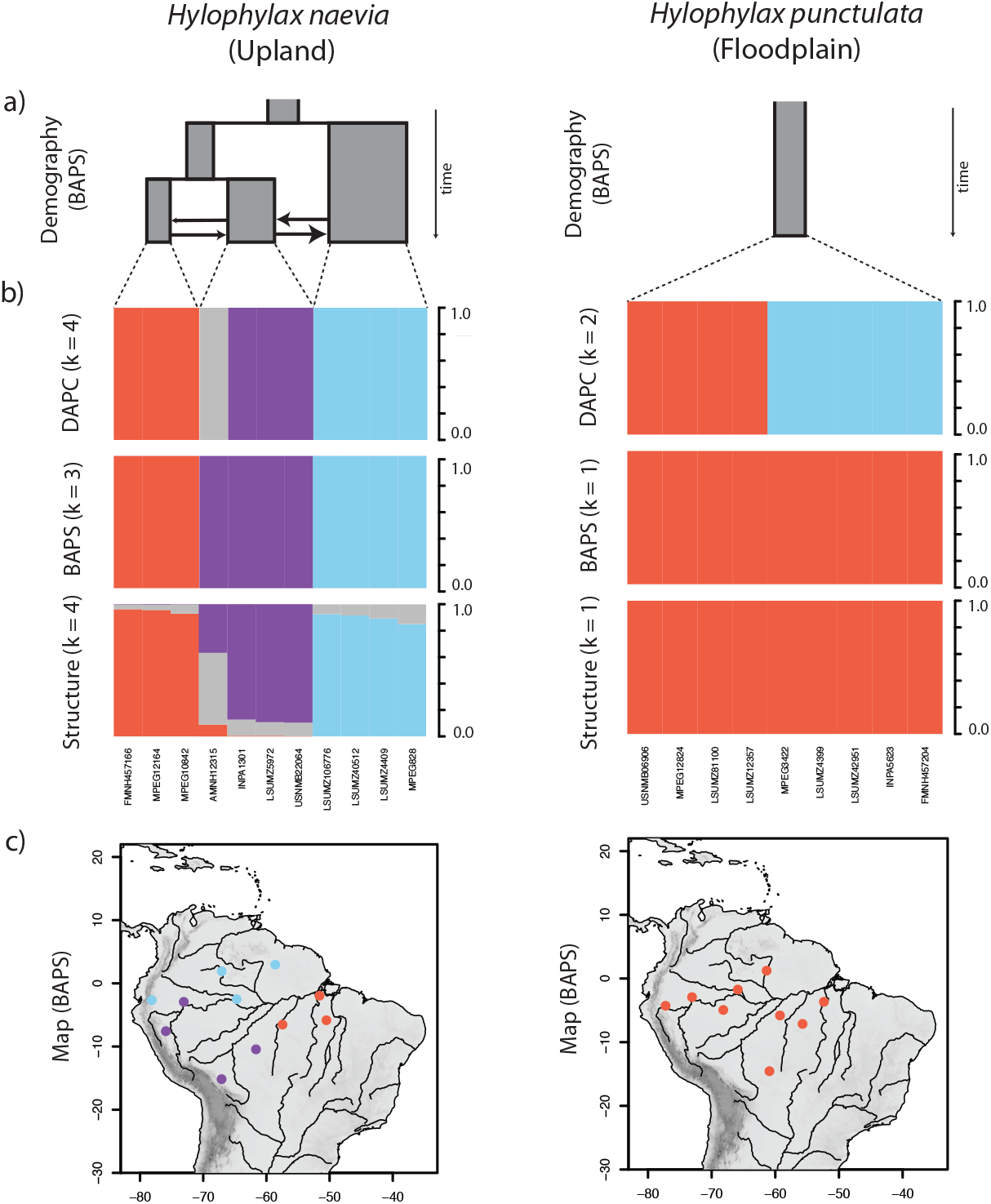
A representative pair of study species depicting (a) demographic models inferred based on the population assignments from BAPS, (b) individual population assignments based on Structure, BAPS, and DAPC analyses, and (c) sample distribution and the distribution of populations inferred with BAPS. The demographic models depict population history through time, with the width of boxes proportional to their mutation-scaled effective population size (*θ*), their depth proportional to relative population divergence times (*τ*), and the size of arrows between them indicating the level of migration between terminal populations.

Genetic metrics compiled for each species from the above analyses and representing dataset attributes, genetic diversity, divergence, and demographic history are presented in table 1 and appendix D. Each genetic metric was correlated (*P* < 0.05) with between 1 and 13 other metrics (mean = 9; fig. A4), and we clustered the variables into 7 groups containing high within-group correlations. Nine of 18 genetic metrics exhibited phylogenetic signal based on Pagel’s λ tests (table A6). The level of overall divergence between the species in a pair, however, was not associated with the degree to which they differed in any genetic variable (table A6). Two metrics, number of mapped reads and number of Structure populations, were correlated with whether a species represented a single species or species complex (table A6).

Habitat association predicted (*P* < 0.05) seven genetic metrics from three semi-independent groups in single-comparison GLMM analyses (fig. 3; table 1). Three measures of species-wide genetic diversity, the number of variable sites, nucleotide diversity (π), and the mutation-scaled effective population size (θ), were higher in upland forest species than floodplain species. Tajima’s *D* was slightly lower in upland forest species than floodplain species, although this was partly driven by one outlier (without Collared Trogon, *Trogon collaris t* = -2.02, *P* = .051). Population divergence across the landscape, measured both by d_XY_ and F_ST_, was higher in upland forest species.

**Figure 3:**
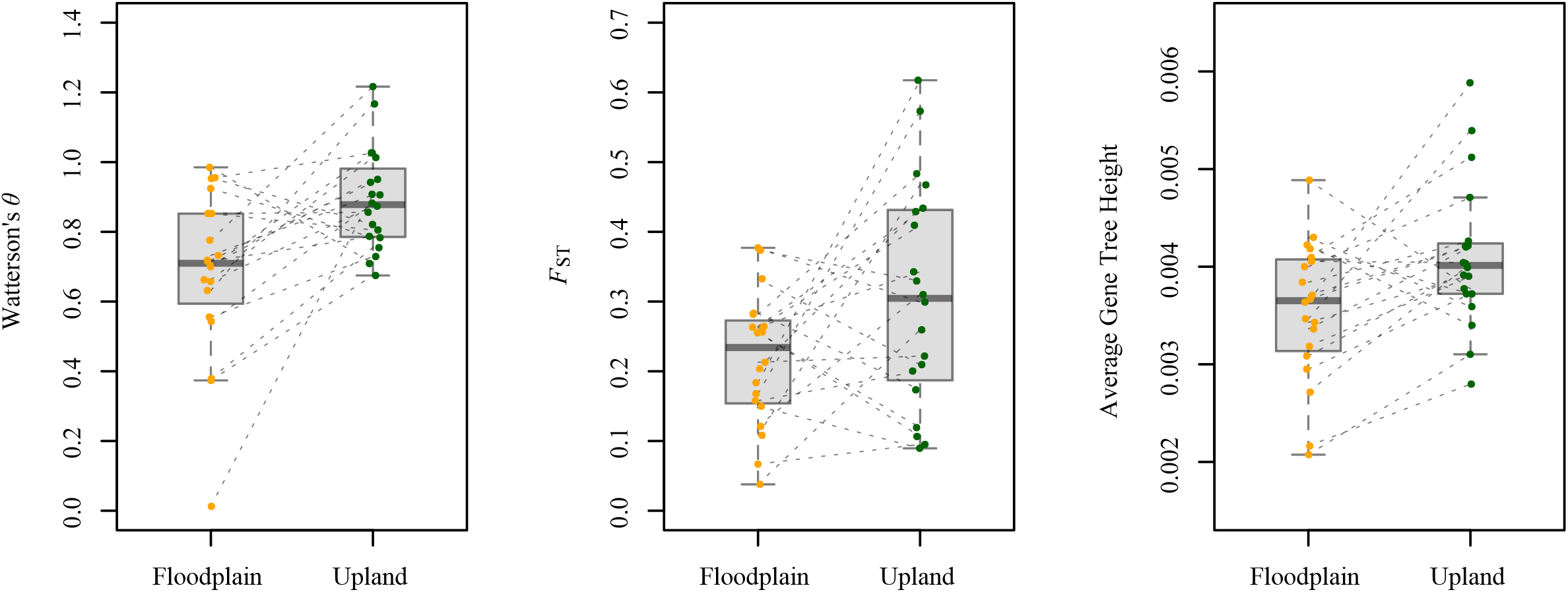
Plots of three representative genetic metrics (each from a different group of correlated variables) that were found to be different between floodplain and upland forest based on GLMM analyses. Dashed lines connect members of a species pair.

Correlations between habitat and d_XY_ or F_ST_ changed little when corrected for small differences among species in the geographic distances between samples (table A7). Finally, the average height of gene trees was greater in upland forest species. PGLS results were similar to those from GLMMs, with greater nucleotide diversity, higher θ, lower Tajima’s *D*, greater gene tree height, and larger d_XY_ and F_ST_ values in canopy than understory species (table 1). In addition, the number of populations inferred using both Structure and DAPC was greater in upland forest species based on PGLS. Relationships changed little in multi-predictor models including forest stratum and/or Kipp’s index and, in genetic metrics that were associated with habitat, *P* values for the other predictor variables were generally non-significant (tables A8-A14). Across response variables, four to seven species pairs showed a difference in the direction opposing the majority of pairs (table A15), with the floodplain species in most of these cases displaying greater diversity or more divergence than the upland forest species.

PGLS analyses with forest stratum or Kipp’s index did not detect strong associations with metrics of population genetic diversity or population history. The only significant relationship was a positive correlation between forest stratum and the relative nucleotide diversity on the Z chromosome versus autosomes (*t* = 2.60, *P* = .01; fig. A5). Results of forest stratum and Kipp’s index comparisons were similar between single- and multi-predictor analyses (tables A8-A14).

## DISCUSSION

We found that the habitat associations of Amazonian birds predict genome-wide estimates of evolutionary metrics representing genetic diversity, divergence across the landscape, number of genetic populations, and depth of gene histories. These results reaffirm the hypothesis that ecological traits of species are useful predictors of intraspecific diversity and evolutionary processes (Loveless and Hamrick 1984; Duminil et al. 2007; Burney and Brumfield 2009; Kisel et al. 2012; Pabijan et al. 2012; Paz et al. 2015). Most of the genetic differences between floodplain and upland species are related to genetic divergence associated with geography. Although species-wide metrics of genetic diversity were associated with habitat, heterozygosity within populations was not, which suggests that differences in genetic diversity among species can be ascribed to variation in metapopulation structure or geographic divergence. Measures of spatial patterns of divergence may be more useful metrics for use in comparative studies than summary estimates of genetic diversity.

We recovered few associations between habitat and historical demographic processes estimated using a multi-population demographic model. This is likely for two reasons. Demographic parameters are notoriously difficult to estimate accurately (Myers et al. 2008; Strasburg and Rieseburg 2010; Schraiber and Akey 2015), and estimates can be spurious when genetic variation is impacted by non-neutral evolutionary processes such as natural selection (Hahn 2008). In addition, we may have low power to detect demographic differences because genetic diversity is partitioned into multiple parameters by demographic models, unlike in most of the genetic diversity and divergence metrics. Detecting concerted trait-based variation in demographic parameters may require better data, improved models, and the development of more sensitive comparative methods for these types of data.

Diverse mechanisms may be responsible for differences in diversity and divergence between floodplain and upland forest species. For example, greater dispersal over either ecological or evolutionary timescales in floodplain species could explain their lower levels of diversity and divergence with respect to upland forest species. Birds of the forest interior are less likely to cross openings than birds of forest edges (Laurance et al. 2004). Seasonal movements are more frequent in birds of edge habitats (Levey and Stiles 1992), and seasonal flooding may annually force some floodplain birds into upland forest, promoting the movement of individuals into new areas (Rosenberg 1990). Rivers, important barriers to dispersal in Amazonia, could be less effective dispersal barriers to floodplain species than to upland species (Capparella 1987; Patton and da Silva 1998). Uplands may not occur within several kilometers of the main channel (Melack and Hess 2011), potentially augmenting the significance of river barriers for upland bird species. River capture events, in which shifts in river courses result in land moving from one bank to the other, may regularly result in the passive movement of patches of floodplain habitat (Salo et al. 1986; Dumont 1991) and associated organisms (Tuomisto and Ruokolainen 1997; Patton et al. 2000) across river barriers, but river capture events involving upland forest may be less frequent (but see Almeida-Filho and Miranda 2007). Kipp’s index did not differ between floodplain and upland forest species, nor did we find higher migration rates in floodplain species than in upland forest species in our demographic models (appendix C). Better metrics of dispersal and gene flow may reveal concerted differences between these habitats that are responsible for differences in genetic divergence.

Differences in population size, population fluctuations through time, or the time a species has been present in the landscape could also explain differences in diversity and divergence between habitats. Floodplains are currently relatively restricted in the Amazon Basin, where they cover about 14% of the lowland area (Melack and Hess 2011). The small area in floodplains may constrain population sizes in floodplain species, leading to lower genetic diversity and fewer opportunities for population divergence. Consistent with this hypothesis, we found lower values of Watterson’s θ, which scales with effective population size, in floodplain species. Differences in climatic or geological history between floodplain and upland areas may also translate into differences in patterns of dispersal and differentiation over long timescales. Sea level rise associated with climatic changes may have reduced the extent of available terrestrial floodplain habitats during the Quaternary Period (Irion et al. 1997), and recent expansion following these or other events could help explain low diversity or divergence in floodplain species (Matocq et al. 2000; Aleixo 2002; Aleixo 2006). There was no evidence for a stronger signal of expansion in floodplains. Low Tajima’s *D* values are expected under recent population expansion, but Tajima’s *D* values were, if anything, higher in floodplain than in upland forest species. There was no difference between floodplains and uplands in the change in population size between the root population and extant populations in G-PhoCS. Recent colonization of the Amazon might also lead to low divergence in floodplains species. The deepest population divergence from G-PhoCS (crown age) is our best metric of the time each species has been distributed in the Amazon Basin. This did not differ, however, between floodplain and upland species.

Overall, the mechanisms causing observed differences in diversity and divergence between floodplain and upland bird species are still unclear. Many processes leave similar signatures in population genetic data (Myers et al. 2008; Strasburg and Rieseburg 2010), which, when combined with uncertainty about environmental history in tropical areas, can make confidently assessing the source of current patterns of diversity challenging (Harvey and Brumfield 2015). More complete and detailed estimates of ecological and evolutionary processes and environmental history may permit better assessments of mechanistic links between habitat preference and genetic diversity and divergence in the future. In the meantime, habitat-associated differences in genetic metrics are interesting in their own right, and for their potential role in longer-term evolutionary dynamics (see below).

Upland forest species, on average, exhibited greater genomic diversity, deeper history, and greater divergence than floodplain species in all significant comparisons. The deep genetic divergences observed in many upland forest species coincided roughly with rivers that represent major putative biogeographic barriers for terrestrial Amazonian species (Cracraft 1985; da Silva et al. 2005). Higher resolution studies are warranted within particular species to better characterize intraspecific diversity and determine whether populations merit recognition as full species. In particular, Variegated Tinamou (*Crypturellus variegatus*), Rufous-capped Antthrush (*Formicarius colma*), Spot-backed Antbird (*Hylophylax naevius*), Sooty Antbird (*Hafferia fortis*), Black-faced Antbird (*Myrmoborus myotherinus*), and Straight-billed Hermit (*Phaethornis bourcieri*) contained deep divergences within currently recognized species. However, some species pairs in all comparisons showed opposing patterns. The species pairs with consistently lower diversity and divergence in the upland forest were *Piaya*, *Formicarius*, *Synallaxis*, and *Saltator*. In *Piaya*, there was no notable divergence within the upland forest species (Black-bellied Cuckoo, *P. melanogaster*), and weak divergence in the floodplain forest species (Squirrel Cuckoo, *P. cayana*). In the last three cases, however, a single deep divergence was present in the floodplain forest species. In Black-faced Antthrush (*Formicarius analis*) and Grayish Saltator (*Saltator coerulescens*), a highly divergent population was present in the Guianan region, whereas in Plain-crowned Spinetail (*Synallaxis gujanensis*) it was in the southwestern Amazon near the foot of the Andes. These species pairs demonstrate that the trend for greater diversity and divergence in upland species is not universal and support the theory that idiosyncrasy is an important component of patterns of intraspecific diversity across Neotropical bird species (Brumfield 2012).

Despite prior evidence that divergence in Neotropical birds is associated with forest stratum or dispersal ability (Burney and Brumfield 2009; Smith et al. 2014b), we detected little evidence for relationships between these traits and genomic diversity. The only relationship recovered between genetic metrics and forest stratum or Kipp’s index involved higher nucleotide diversity on the Z chromosome relative to autosomes in understory species. This is surely due in part to low power resulting from our study design. Members of a genus in our study always exhibited the same forest stratum preference and similar Kipp’s index values, and therefore the effective sample size for these comparisons was roughly half that of the paired comparisons involving habitat. We expect that improved sample sizes would find stronger associations between forest stratum and genetic metrics. The forest canopy is in many ways analogous to edge habitats like those found in floodplains, and both are thought to harbor higher concentrations of birds that undergo seasonal movements than the interior of tall forest (Levey and Stiles 1992). Both canopy and floodplain bird species have lower subspecies richness than understory and upland forest species, respectively (Salisbury et al. 2012).

We have demonstrated that an ecological trait, habitat associations, predicts variation across species in population genetics and evolutionary metrics. Birds in the interior of upland forest have greater diversity, more divergence across the landscape, and deeper gene histories than birds of floodplain and edge habitats. Habitat-associated differences in levels and patterns of variation are significant because they may reflect different propensities to respond to environmental change, form new species, and succumb to extinction. Interestingly, the upland forest avifauna is more diverse (1,058 species) than the floodplain forest avifauna (154 species) in the Amazon Basin (Parker et al. 1996). Because population divergence and speciation rates over long evolutionary timescales may show similar associations with ecological traits (Riginos et al. 2014), different rates of population divergence between upland forest and floodplains may have played a role in producing their disparate diversities via species selection (Stanley 1975). Different conservation strategies may also be necessary to preserve the divergent patterns of genetic diversity and evolutionary processes observed in upland and floodplain regions. Practically, we have demonstrated that genomic datasets can be used to estimate diverse parameters for testing hypotheses about traits associated with genomic diversity. Studies examining additional taxa and new methods for estimating more detailed population histories are sure to provide more insight into the impacts of ecology on population genomics and evolution in the near future. In addition, more detailed ecological information will be needed to permit better quantification of ecological traits for comparison with evolutionary variables.

## Acknowledgements

We thank the field workers and museum curators and staff that obtained material and maintain the specimens and genetic resources used in this study. Special thanks to the American Museum of Natural History, Instituto Nacional de Pesquisas da Amazônia, Louisiana State University Museum of Natural Science, Museu Paraense Emilio Goeldi, Museu de Zoologia at the Universidade de São Paulo, Field Museum of Natural History, National Museum of Natural History (Smithsonian Institution), and University of Kansas Natural History Museum. D. F. Lane, B. M. Whitney, J. V. Remsen Jr., and L. N. Naka helped with study design and species selection. J. M. Brown, M. E. Hellberg, J. V. Remsen Jr., B. M. Winger, S. Singhal, L. L. Knowles, the Brumfield and Rabosky labs, and the LSUMNS Vert Lunch group provided helpful discussion. Funding was provided by NSF grants DEB-1210556 (to M.G.H. and R.T.B.) and DEB-1146265 (to R.T.B.), the Nuttall Ornithological Club, and the American Ornithologist’s Union. A.A. and C.C.R. are supported by Brazilian Research Council (CNPq) productivity fellowships.

## Appendix A From “Habitat preference predicts genetic diversity and population divergence in Amazonian birds”

### Supplementary Figures and Tables

**Figure A1:**
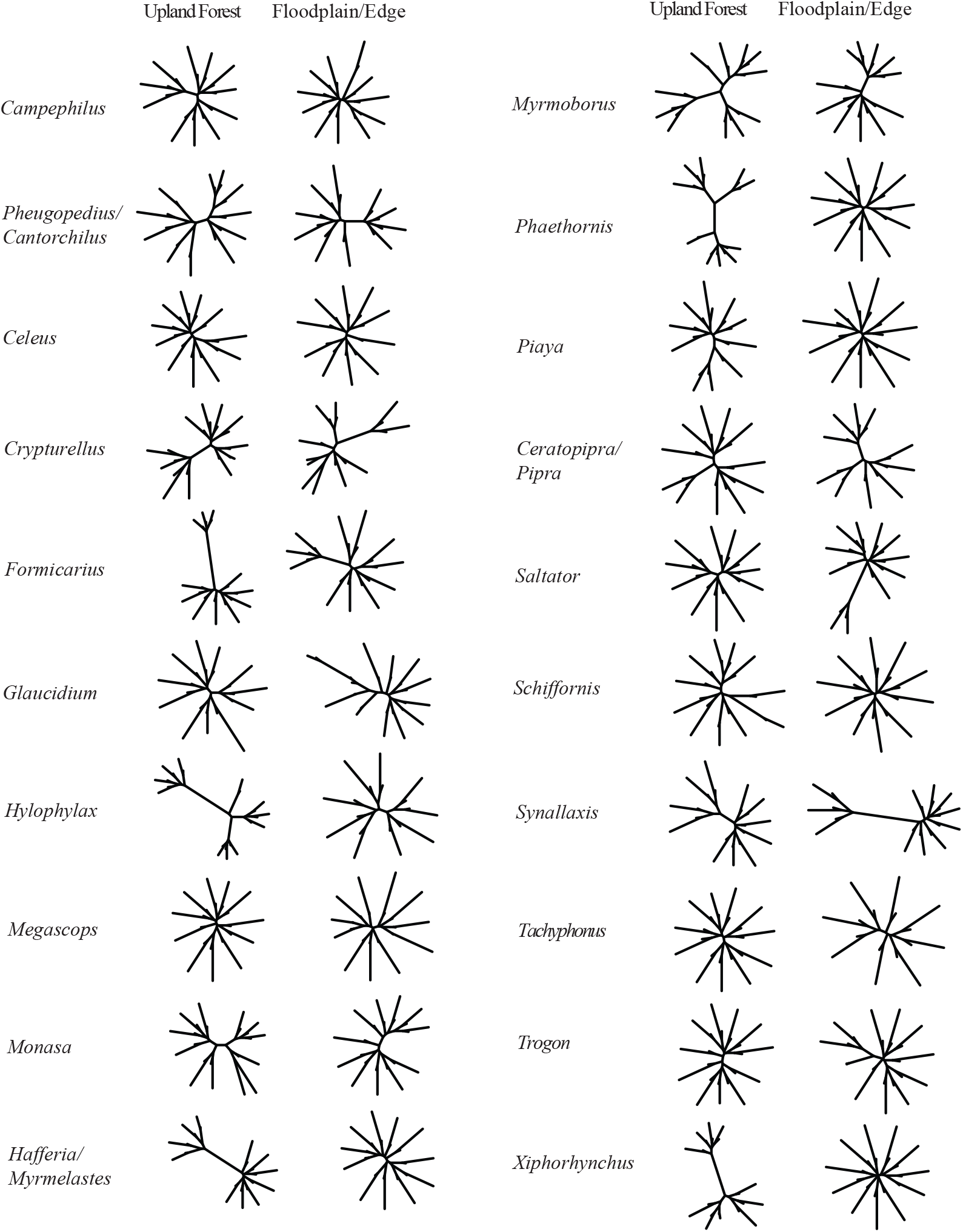
Unrooted MrBayes trees of concatenated SNPs from both alleles in each individual after outgroup, mis-identified, contaminated, and failed samples were removed. In each individual, the rarer allele in the population was generally assigned to the second haplotype. As a result, many individuals are represented by one short and one long terminal branch.

**Figure A2:**
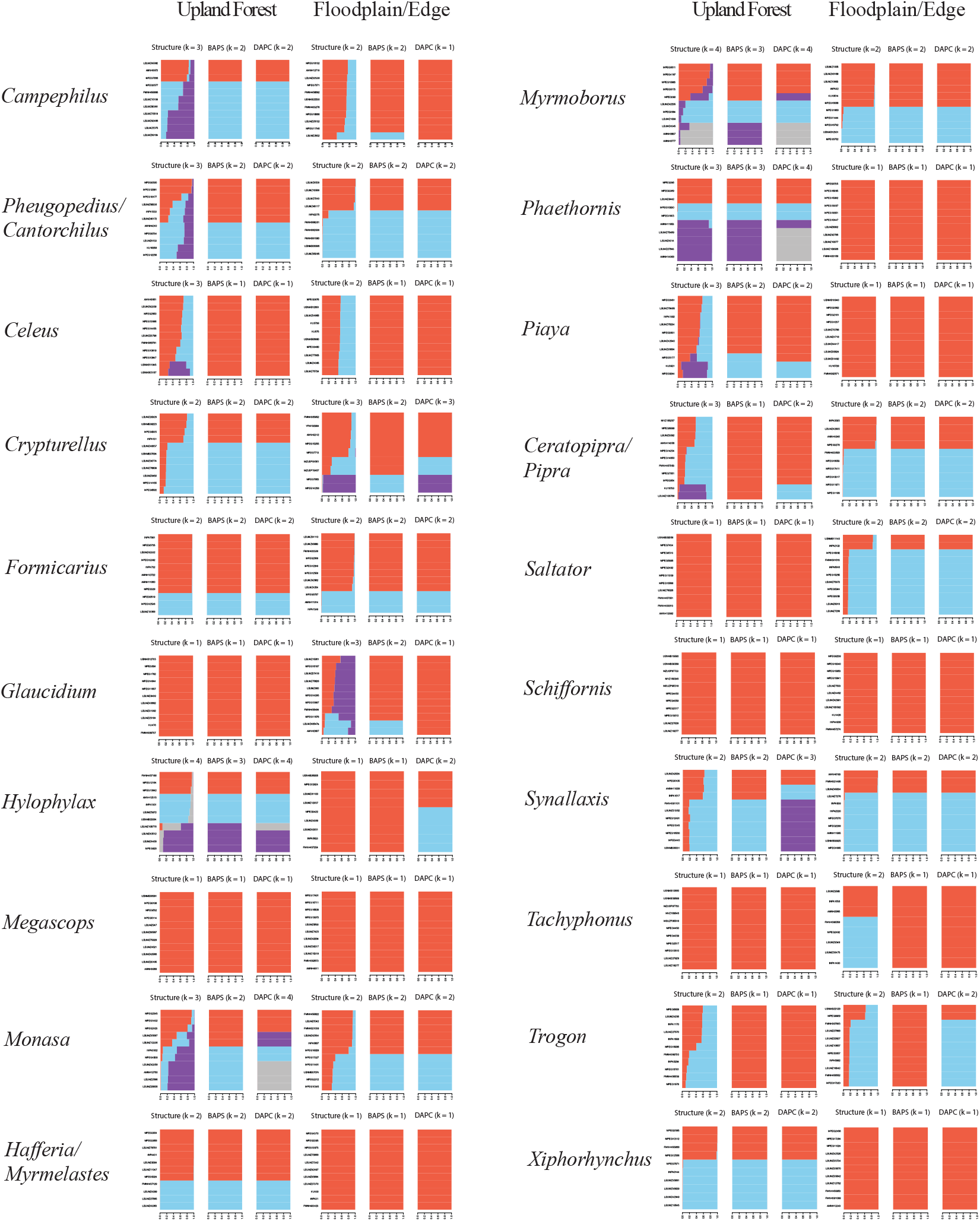
Plots of population genetic structure and cluster assignments inferred from Structure, BAPS, and DAPC for all 40 study species. Distinct colors represent different clusters, and the size of bars is proportional to the probability of assignment to a particular cluster.

**Figure A3:**
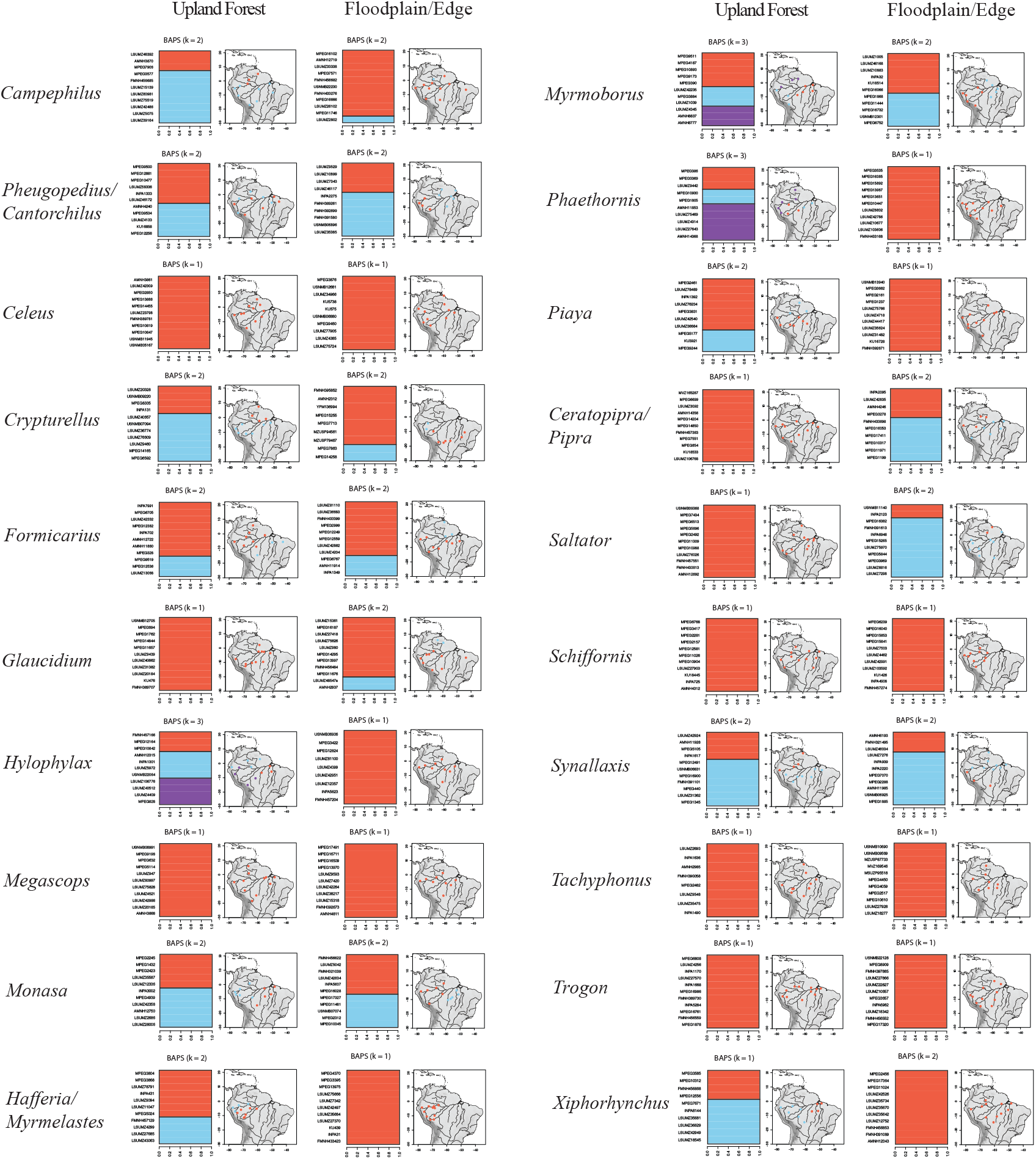
Population genetic structure and cluster assignments from BAPS adjoining maps showing the distribution of samples assigned to each cluster across the Amazon Basin.

**Figure A4:**
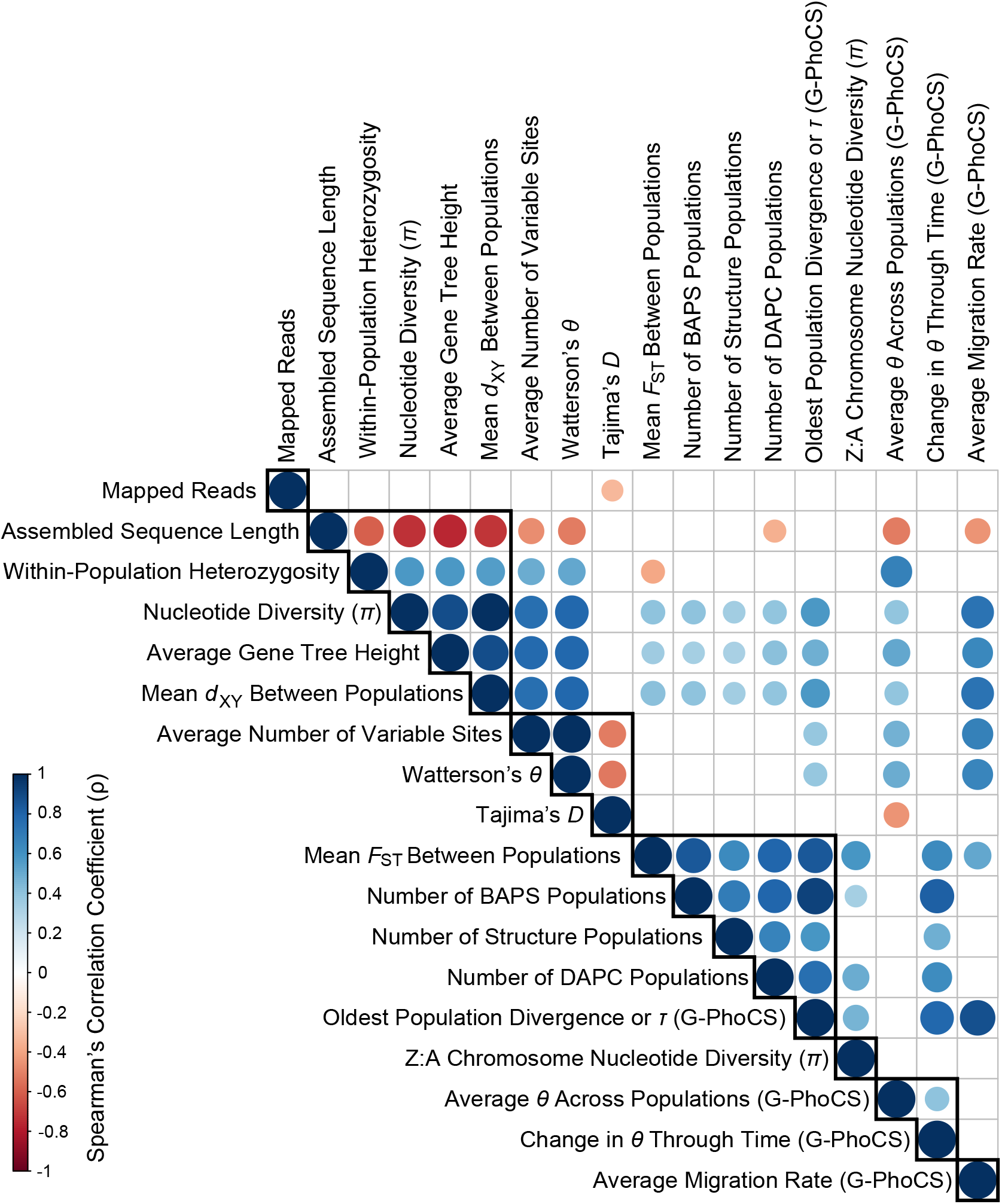
Pairwise Spearman’s correlation coefficients between all genetic metrics. The size of circles reflects relative significance levels, and empty cells represent no significant correlation (*P* > 0.05). Black polygons outline seven groups of highly correlated variables based on ClustOfVar analysis.

**Figure A5:**
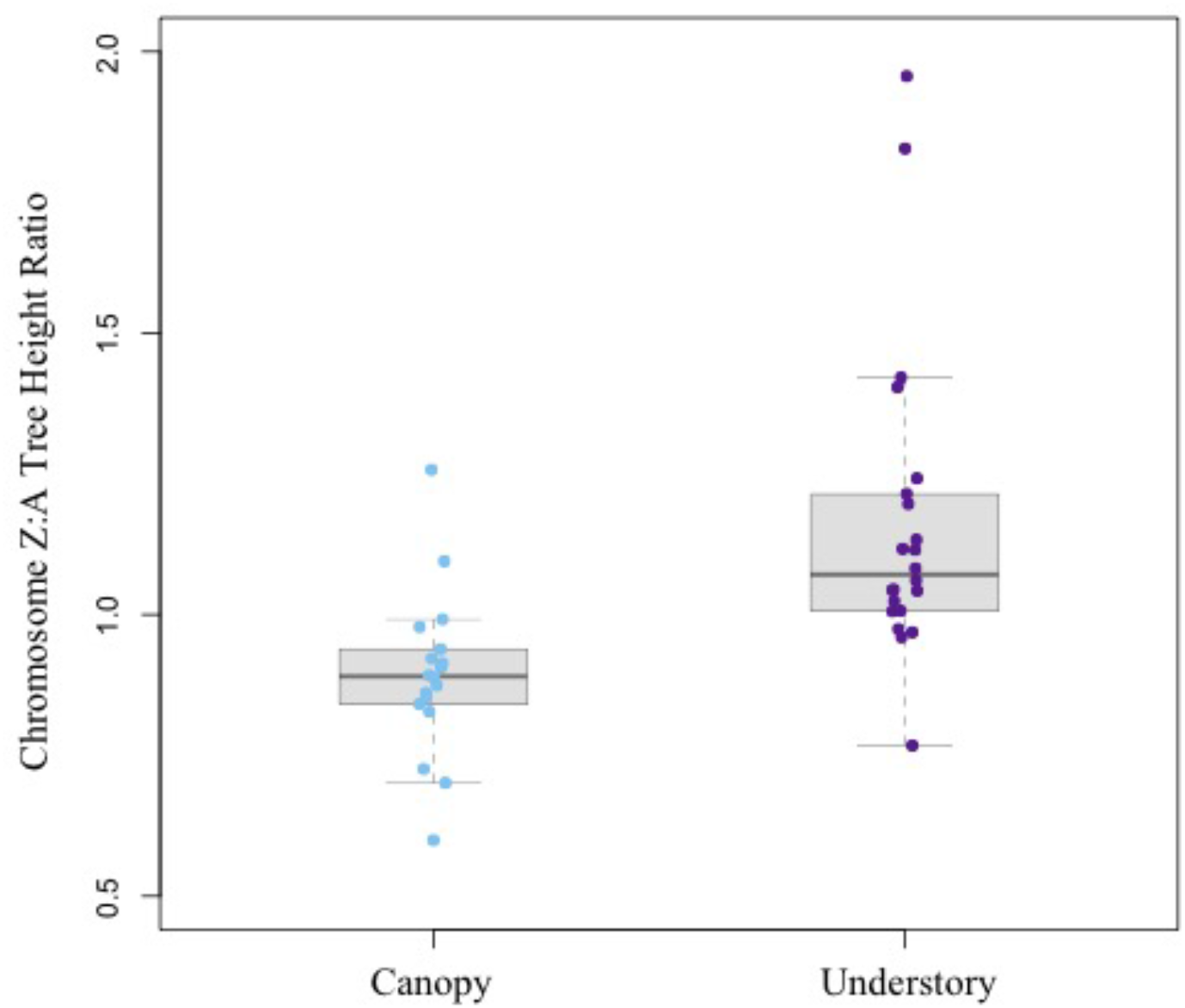
A plot showing the ratio of nucleotide diversity between loci mapping to the Z chromosome and those mapping to the autosomes versus forest stratum. The outliers in the understory group represent the two *Crypturellus* tinamous.

**Table A1:** The mitochondrial genome used for reference-based assembly of partial mitochondrial genomes in each pair of study species

| Study Species | Reference Mitochondrial Genome | Genbank # |
| --- | --- | --- |
| *Campephilus melanoleucos* | *Dryocopus pileatus* | DQ780879 |
| *Campephilus rubricollis* | *Dryocopus pileatus* | DQ780879 |
| *Cantorchilus leucotis* | *Henicorhina leucosticta* | KJ719074 |
| *Celeus flavus* | *Dryocopus pileatus* | DQ780879 |
| *Celeus grammicus/undatus* | *Dryocopus pileatus* | DQ780879 |
| *Crypturellus undulatus* | *Tinamus guttatus* | KR149454 |
| *Crypturellus variegatus* | *Tinamus guttatus* | KR149454 |
| *Formicarius analis* | *Scytalopus magellanicus* | KJ909189 |
| *Formicarius colma* | *Scytalopus magellanicus* | KJ909189 |
| *Glaucidium brasilianum* | *Ninox novaeseelandiae* | AY309457 |
| *Glaucidium hardyi* | *Ninox novaeseelandiae* | AY309457 |
| *Hylophylax naevius* | *Thamnophilus nigrocinereus* | KJ909192 |
| *Hylophylax punctulatus* | *Thamnophilus nigrocinereus* | KJ909192 |
| *Megascops choliba* | *Strix leptogrammica* | KC953095 |
| *Megascops watsonii* | *Strix leptogrammica* | KC953095 |
| *Monasa morphoeus* | *Dryocopus pileatus* | DQ780879 |
| *Monasa nigrifrons* | *Dryocopus pileatus* | DQ780879 |
| *Hafferia fortis* | *Thamnophilus nigrocinereus* | KJ909192 |
| *Myrmelastes hyperythra* | *Thamnophilus nigrocinereus* | KJ909192 |
| *Myrmoborus leucophrys* | *Thamnophilus nigrocinereus* | KJ909192 |
| *Myrmoborus myotherinus* | *Thamnophilus nigrocinereus* | KJ909192 |
| *Phaethornis bourcieri/philippii* | *Amazilia versicolor* | KF624601 |
| *Phaethornis hispidus* | *Amazilia versicolor* | KF624601 |
| *Pheugopedius coraya/genibarbis* | *Henicorhina leucosticta* | KJ719074 |
| *Piaya cayana* | *Eudynamys taitensis* | EU410487 |
| *Piaya melanogaster* | *Eudynamys taitensis* | EU410487 |
| *Pipra aureola/fasciicauda/filicauda* | *Lepidothrix coronata* | KJ909196 |
| *Ceratopipra chloromeros/erythrocephala/rubrocapilla* | *Lepidothrix coronata* | KJ909196 |
| *Saltator coerulescens* | *Thraupis episcopus* | KM078765 |
| *Saltator grossus* | *Thraupis episcopus* | KM078765 |
| *Schiffornis major* | *Cnemotriccus fuscatus* | AY596278 |
| *Schiffornis turdina* | *Cnemotriccus fuscatus* | AY596278 |
| *Synallaxis gujanensis* | *Scytalopus magellanicus* | KJ909189 |
| *Synallaxis rutilans* | *Scytalopus magellanicus* | KJ909189 |
| *Tachyphonus cristatus* | *Thraupis episcopus* | KM078765 |
| *Tachyphonus luctuosus* | *Thraupis episcopus* | KM078765 |
| *Trogon collaris* | *Trogon viridis* | EU410490 |
| *Trogon rufus* | *Trogon viridis* | EU410490 |
| *Xiphorhynchus elegans/spixii* | *Scytalopus magellanicus* | KJ909189 |
| *Xiphorhynchus obsoletus* | *Scytalopus magellanicus* | KJ909189 |

**Table A2:**
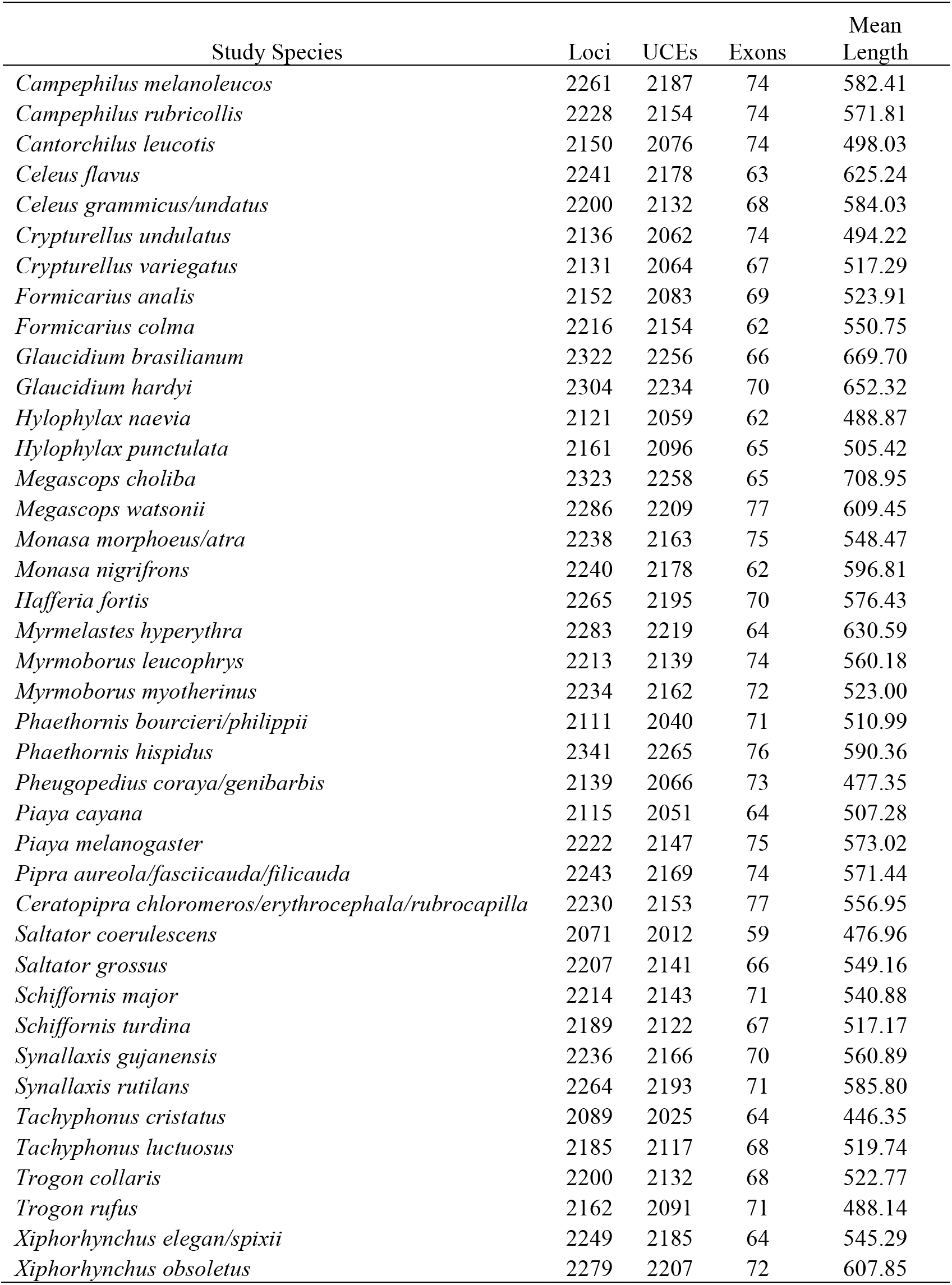
Summary information on loci recovered for each species

| Study Species | Loci | UCEs | Exons | Mean Length |
| --- | --- | --- | --- | --- |
| *Campephilus melanoleucos* | 2261 | 2187 | 74 | 582.41 |
| *Campephilus rubricollis* | 2228 | 2154 | 74 | 571.81 |
| *Cantorchilus leucotis* | 2150 | 2076 | 74 | 498.03 |
| *Celeus flavus* | 2241 | 2178 | 63 | 625.24 |
| *Celeus grammicus/undatus* | 2200 | 2132 | 68 | 584.03 |
| *Crypturellus undulatus* | 2136 | 2062 | 74 | 494.22 |
| *Crypturellus variegatus* | 2131 | 2064 | 67 | 517.29 |
| *Formicarius analis* | 2152 | 2083 | 69 | 523.91 |
| *Formicarius colma* | 2216 | 2154 | 62 | 550.75 |
| *Glaucidium brasilianum* | 2322 | 2256 | 66 | 669.70 |
| *Glaucidium hardyi* | 2304 | 2234 | 70 | 652.32 |
| *Hylophylax naevia* | 2121 | 2059 | 62 | 488.87 |
| *Hylophylax punctulata* | 2161 | 2096 | 65 | 505.42 |
| *Megascops choliba* | 2323 | 2258 | 65 | 708.95 |
| *Megascops watsonii* | 2286 | 2209 | 77 | 609.45 |
| *Monasa morphoeus/atra* | 2238 | 2163 | 75 | 548.47 |
| *Monasa nigrifrons* | 2240 | 2178 | 62 | 596.81 |
| *Hafferia fortis* | 2265 | 2195 | 70 | 576.43 |
| *Myrmelastes hyperythra* | 2283 | 2219 | 64 | 630.59 |
| *Myrmoborus leucophrys* | 2213 | 2139 | 74 | 560.18 |
| *Myrmoborus myotherinus* | 2234 | 2162 | 72 | 523.00 |
| *Phaethornis bourcieri/philippii* | 2111 | 2040 | 71 | 510.99 |
| *Phaethornis hispidus* | 2341 | 2265 | 76 | 590.36 |
| *Pheugopedius coraya/genibarbis* | 2139 | 2066 | 73 | 477.35 |
| *Piaya cayana* | 2115 | 2051 | 64 | 507.28 |
| *Piaya melanogaster* | 2222 | 2147 | 75 | 573.02 |
| *Pipra aureola/fasciicauda/filicauda* | 2243 | 2169 | 74 | 571.44 |
| *Ceratopipra chloromeros/erythrocephala/rubrocapilla* | 2230 | 2153 | 77 | 556.95 |
| *Saltator coerulescens* | 2071 | 2012 | 59 | 476.96 |
| *Saltator grossus* | 2207 | 2141 | 66 | 549.16 |
| *Schiffornis major* | 2214 | 2143 | 71 | 540.88 |
| *Schiffornis turdina* | 2189 | 2122 | 67 | 517.17 |
| *Synallaxis gujanensis* | 2236 | 2166 | 70 | 560.89 |
| *Synallaxis rutilans* | 2264 | 2193 | 71 | 585.80 |
| *Tachyphonus cristatus* | 2089 | 2025 | 64 | 446.35 |
| *Tachyphonus luctuosus* | 2185 | 2117 | 68 | 519.74 |
| *Trogon collaris* | 2200 | 2132 | 68 | 522.77 |
| *Trogon rufus* | 2162 | 2091 | 71 | 488.14 |
| *Xiphorhynchus elegans/spixii* | 2249 | 2185 | 64 | 545.29 |
| *Xiphorhynchus obsoletus* | 2279 | 2207 | 72 | 607.85 |

**Table A3:**
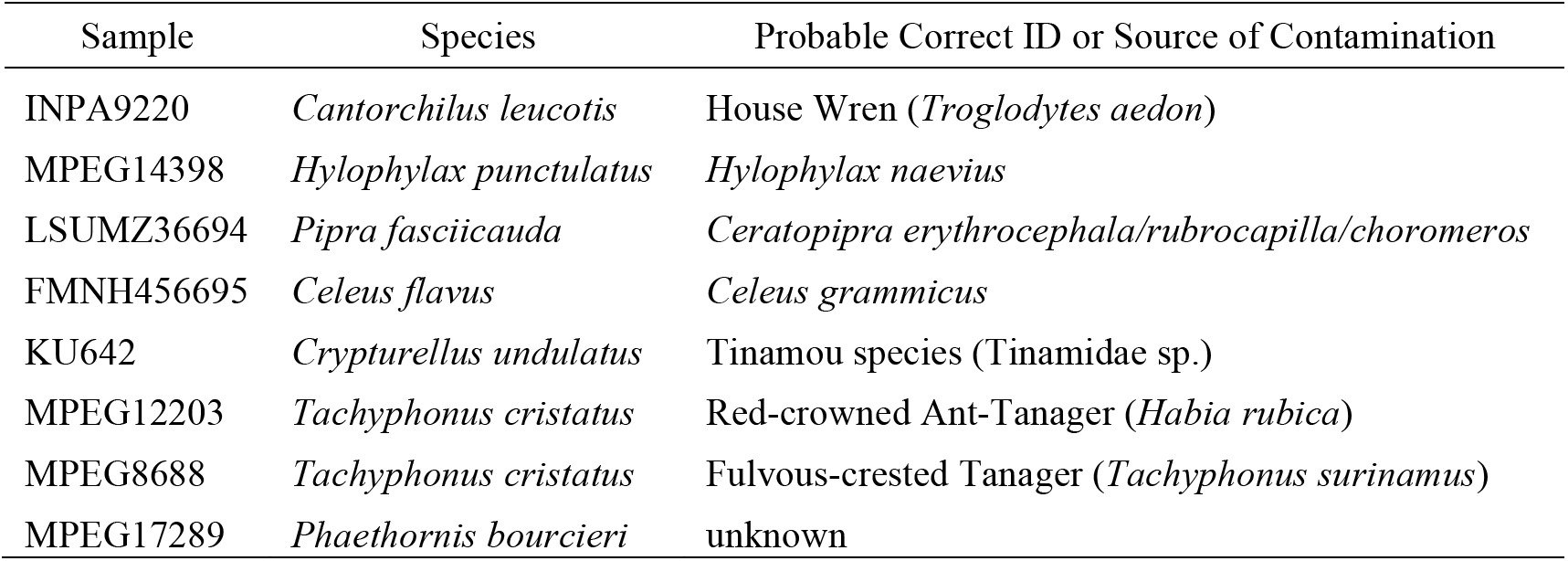
Samples that were likely mis-identified or heavily contaminated with another sample

**Table A4:**
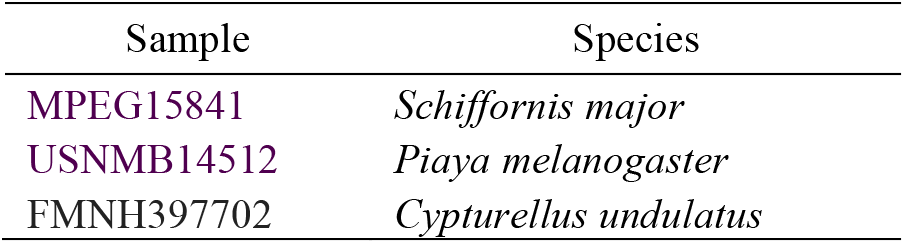
Samples that included DNA from multiple sources or were subject to high error rates based on an excess of single-copy SNP alleles

**Table A5:** Samples that were removed from analysis due to a high frequency of missing data at variable sites

| Sample | Species | % Missing Data |
| --- | --- | --- |
| MPEG12248 | *Hylophylax punctulatus* | 99 |
| MPEG1439 | *Tachyphonus cristatus* | 92 |
| FMNH397598 | *Xiphorhynchus spixii* | 85 |

**Table A6:** Results of tests for phylogenetic signal in each genetic metric, PGLS comparing difference in each metric within a pair to their overall divergence, and PGLS analysis of each metric versus whether species are considered one or multiple taxonomic species

| Variable | Phylogenetic Signal |  | Correlation with Divergence Between Species Pair |  | Correlation with Whether Species or Species Complex |  |
| --- | --- | --- | --- | --- | --- | --- |
| | $\lambda$ | $P$ | $R^2$ | $P$ | $R^2$ | $P$ |
| Mapped Reads | .16 | .41 | -.004 | .35 | .13 | .01 |
| Assembled Sequence Length | .91 | <.001 | -.05 | .80 | -.02 | .73 |
| Average Number of Variable Sites | .53 | .14 | -.04 | .63 | -.001 | .34 |
| Nucleotide Diversity ($\pi$) | .75 | <.001 | -.05 | .78 | -.02 | .57 |
| Z:A Chromosome Nucleotide Diversity ($\pi$) Ratio | .90 | <.001 | -.05 | .68 | -.02 | .52 |
| Watterson's $\theta$ | .53 | .11 | -.05 | .75 | .00 | .32 |
| Tajima's $D$ | .53 | .11 | .01 | .28 | -.006 | .39 |
| Within-Population Heterozygosity | .95 | <.001 | -0.06 | .97 | -.02 | .56 |
| Average Gene Tree Height | .85 | <.001 | -.05 | .87 | -.02 | .57 |
| Mean $d_{XY}$ Between Populations | .73 | <.001 | -.05 | .77 | -.02 | .55 |
| Mean $F_{ST}$ Between Populations | .29 | .16 | .006 | .30 | .04 | .10 |
| Number of Structure Populations | .05 | .82 | -.03 | .50 | .09 | .03 |
| Number of BAPS Populations | .19 | .21 | -.05 | .72 | .04 | .12 |
| Number of DAPC Populations | .49 | .03 | -.04 | .59 | -.01 | .51 |
| Average $\theta$ Across Populations (G-PhoCS) | .85 | <.001 | -.05 | .78 | -.02 | .72 |
| Change in $\theta$ Through Time (G-PhoCS) | .00 | 1.00 | .02 | .26 | -.02 | .55 |
| Oldest Population Divergence or $\tau$ (G-PhoCS) | .40 | .006 | -.01 | .41 | -.02 | .53 |
| Average Migration Rate (G-PhoCS) | .02 | .91 | -.15 | .68 | -.03 | .51 |

**Table A7:** Results of GLMMs and PGLS of habitat versus F_ST_ and d_XY_ controlling for differences among species in geographic distance between samples

| Variable | GLMM $t$ | GLMM $P$ | PGLS $t$ | PGLS $P$ |
| --- | --- | --- | --- | --- |
| Mean $d_{XY}$ Between Populations / Distance | 1.87 | .07 | 3.642 | <.001 |
| Mean $F_{ST}$ Between Populations / Distance | 2.15 | .04 | 3.05 | .004 |

**Table A8:**
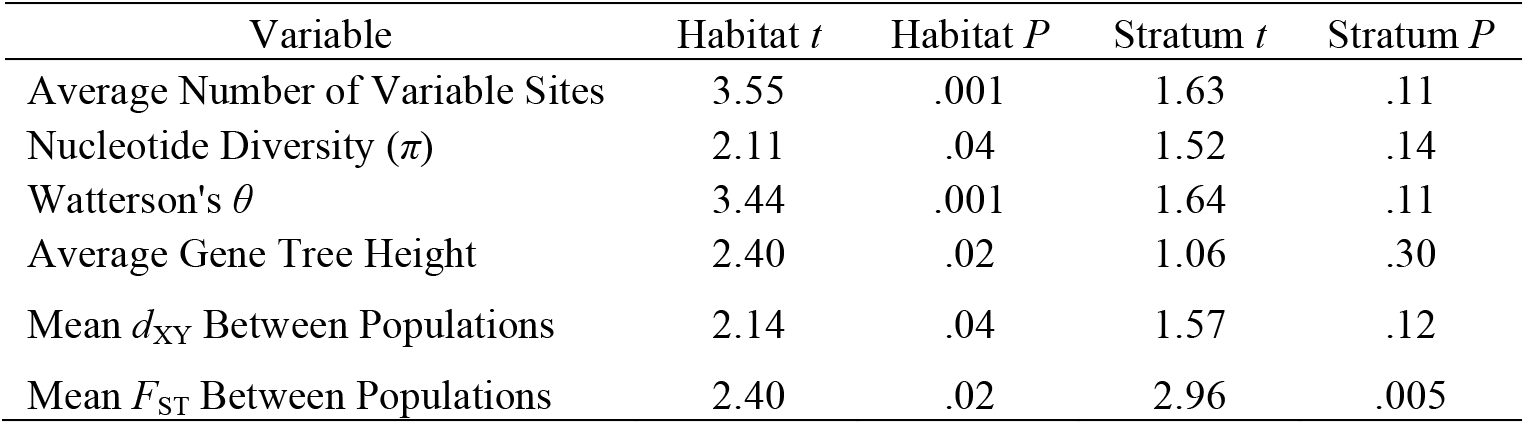
Significant results from GLMMs with habitat based on multi-predictor analysis with forest stratum included

**Table A9:** Significant results from GLMMs with habitat based on multi-predictor analysis with Kipp’s index included

| Variable | Habitat $t$ | Habitat $P$ | Kipp's $t$ | Kipp's $P$ |
| --- | --- | --- | --- | --- |
| Average Number of Variable Sites | 3.51 | .001 | -1.42 | .16 |
| Nucleotide Diversity ($\pi$) | 2.06 | .047 | -.66 | .52 |
| Watterson's $\theta$ | 3.40 | .002 | -1.40 | .17 |
| Tajima's $D$ | -2.06 | .050 | 1.54 | .13 |
| Average Gene Tree Height | 2.37 | .023 | -.62 | .54 |
| Mean $d_{XY}$ Between Populations | 2.07 | .045 | -.65 | .52 |
| Mean $F_{ST}$ Between Populations | 2.16 | .04 | -.11 | .91 |

**Table A10:** Significant results from GLMMs with habitat based on multi-predictor analysis with forest stratum and Kipp’s index included

| Variable | Habitat $t$ | Habitat $P$ | Kipp's $t$ | Kipp's $P$ | Stratum $t$ | Stratum $P$ |
| --- | --- | --- | --- | --- | --- | --- |
| Average Number of Variable Sites | 3.58 | .001 | -1.35 | .18 | 1.56 | .13 |
| Nucleotide Diversity ($\pi$) | 2.09 | .04 | -.58 | .56 | 1.47 | .15 |
| Watterson's $\theta$ | 3.46 | .001 | -1.34 | .19 | 1.58 | .12 |
| Tajima's $D$ | -2.03 | .050 | 1.5 | .14 | -.34 | .74 |
| Average Gene Tree Height | 2.37 | .02 | -.56 | .58 | 1.02 | .32 |
| Mean $d_{XY}$ Between Populations | 2.11 | .04 | -.57 | .57 | 1.52 | .14 |
| Mean $F_{ST}$ Between Populations | 2.37 | .023 | .048 | .96 | 2.91 | .006 |

**Table A11:** Significant results from PGLS with habitat and forest stratum as predictor

| Variable | Habitat $t$ | Habitat $P$ | Stratum $t$ | Stratum $P$ |
| --- | --- | --- | --- | --- |
| Average Number of Variable Sites | 4.88 | <.001 | .97 | .34 |
| Nucleotide Diversity ($\pi$) | 3.24 | .003 | .60 | .55 |
| Z:A Chromosome Nucleotide Diversity ($\pi$) Ratio | 1.07 | .29 | 2.56 | .01 |
| Watterson's $\theta$ | 4.77 | <.001 | .92 | .36 |
| Tajima's $D$ | -2.27 | .03 | -.28 | .78 |
| Average Gene Tree Height | 3.54 | .001 | .55 | .58 |
| Mean $d_{XY}$ Between Populations | 3.25 | .002 | .61 | .55 |
| Mean $F_{ST}$ Between Populations | 2.84 | .007 | .51 | .62 |
| Number of Structure Populations | 2.78 | .008 | -.35 | .73 |
| Number of DAPC Populations | 2.83 | .007 | .86 | .40 |

**Table A12:** Significant results from PGLS with habitat and Kipp’s index as predictor variables

| Variable | Habitat $t$ | Habitat $P$ | Kipp's $t$ | Kipp's $P$ |
| --- | --- | --- | --- | --- |
| Average Number of Variable Sites | -1.38 | .17 | 4.83 | <.001 |
| Nucleotide Diversity ($\pi$) | 3.21 | .003 | -.32 | .75 |
| Watterson's $\theta$ | -1.32 | .19 | 4.71 | <.001 |
| Tajima's $D$ | -2.20 | .03 | 1.12 | .27 |
| Average Gene Tree Height | 3.50 | .001 | -.32 | .75 |
| Mean $d_{XY}$ Between Populations | 3.21 | .003 | -.31 | .76 |
| Mean $F_{ST}$ Between Populations | 2.81 | .008 | -.39 | .70 |
| Number of Structure Populations | 2.71 | .01 | -.54 | .59 |
| Number of DAPC Populations | 2.82 | .008 | -.045 | .96 |

**Table A13:** Significant results from PGLS with forest stratum and Kipp’s index as predictor variables

| Variable | Kipp's $t$ | Kipp's $P$ | Stratum $t$ | Stratum $P$ |
| --- | --- | --- | --- | --- |
| Z:A Chromosome Nucleotide Diversity ($\pi$) Ratio | .59 | .56 | 2.61 | .01 |

**Table A14:**
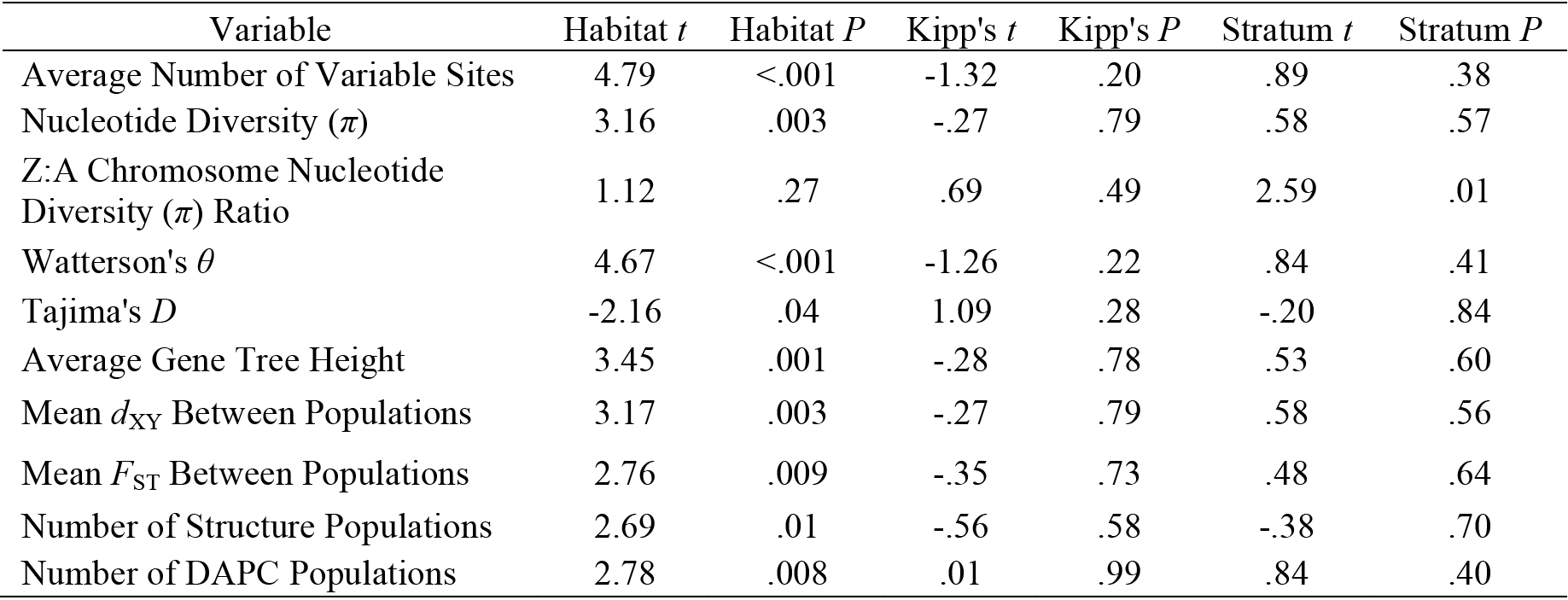
Significant results from PGLS with habitat, forest stratum, and Kipp’s index as predictor variables

| Variable | Habitat $t$ | Habitat $P$ | Kipp's $t$ | Kipp's $P$ | Stratum $t$ | Stratum $P$ |
| --- | --- | --- | --- | --- | --- | --- |
| Average Number of Variable Sites | 4.79 | <.001 | -1.32 | .20 | .89 | .38 |
| Nucleotide Diversity ($\pi$) | 3.16 | .003 | -.27 | .79 | .58 | .57 |
| Z:A Chromosome Nucleotide Diversity ($\pi$) Ratio | 1.12 | .27 | .69 | .49 | 2.59 | .01 |
| Watterson's $\theta$ | 4.67 | <.001 | -1.26 | .22 | .84 | .41 |
| Tajima's $D$ | -2.16 | .04 | 1.09 | .28 | -.20 | .84 |
| Average Gene Tree Height | 3.45 | .001 | -.28 | .78 | .53 | .60 |
| Mean $d_{XY}$ Between Populations | 3.17 | .003 | -.27 | .79 | .58 | .56 |
| Mean $F_{ST}$ Between Populations | 2.76 | .009 | -.35 | .73 | .48 | .64 |
| Number of Structure Populations | 2.69 | .01 | -.56 | .58 | -.38 | .70 |
| Number of DAPC Populations | 2.78 | .008 | .01 | .99 | .84 | .40 |

**Table A15:** Species pairs in which the difference in a genetic metric was opposite that of the overall difference

| Species Pair | Average<br>Number of<br>Variable<br>Sites | Nucleotide<br>Diversity ($\pi$) | Watterson's $\theta$ | Tajima's<br>$D$ | Average<br>Gene Tree<br>Height | Mean $d_{XY}$<br>Between<br>Populations | Mean $F_{ST}$<br>Between<br>Populations |
| --- | --- | --- | --- | --- | --- | --- | --- |
| *Crypturellus* |  |  |  |  |  |  | X |
| *Piaya* | X | X | X | X | X | X |  |
| *Glaucidium* |  |  |  |  |  |  | X |
| *Megascops* |  |  |  |  |  |  | X |
| *Phaethornis* | X |  |  | X |  |  |  |
| *Trogon* |  |  |  |  | X |  | X |
| *Monasa* |  | X |  |  |  |  |  |
| *Hylophylax* |  |  |  | X |  |  |  |
| *Hafferia/Myrmelastes* |  |  |  | X |  |  |  |
| *Myrmoborus* |  |  |  | X |  |  |  |
| *Formicarius* | X | X | X | X | X | X |  |
| *Pipra/Ceratopipra* |  |  |  |  |  |  | X |
| *Synallaxis* | X | X | X |  | X | X | X |
| *Saltator* | X | X | X |  | X | X | X |

